## Supplementary Material for "Cortical connectivity, local dynamics and stability correlates of global conscious states"

### 1 Supplementary Note

#### 1.1 Definition of matrices A, B, and C in neural mass model

The structure of the matrix **A** is composed of independent diagonal blocks, arranged as follows

$$\mathbf{A} = \begin{bmatrix} \mathbf{\Psi} & \mathbf{0} \\ \mathbf{0} & \mathbf{I}_{n_\theta, n_\theta} \end{bmatrix}. \tag{1}$$

Here, $\mathbf{I}_{n_\theta,n_\theta} \in \mathbb{R}^{n_\theta \times n_\theta}$ represents the identity matrix, where $n_\theta$ corresponds to the total number of parameters within the system. The component $\mathbf{\Psi} \in \mathbb{R}^{n_x \times n_x}$ is another diagonal matrix formed from smaller sub-blocks, with $n_x$ denoting the state count

$$\mathbf{\Psi} = \mathrm{diag}(\mathbf{\Psi}_j). \tag{2}$$

Each sub-matrix $\mathbf{\Psi}_j$ in $\mathbf{\Psi}$ is expressed as

$$\mathbf{\Psi}_j = \begin{bmatrix} 0 & 1 \\ -\frac{1}{\tau_j^2} & -\frac{2}{\tau_j} \end{bmatrix}, \tag{3}$$

where $j$ iterates over the range $1, 2, \ldots, N$, representing different connections.

For a discrete representation, $\mathbf{A}_\delta$ is derived from $\mathbf{A}$ as follows

$$\mathbf{A}_\delta = \begin{bmatrix} \mathbf{I} + \delta\mathbf{\Psi} & \mathbf{0} \\ \mathbf{0} & \mathbf{I} \end{bmatrix}. \tag{4}$$

The matrix **B** is configured with the layout

$$\mathbf{B} = \begin{bmatrix} \mathbf{0}_{n_x,n_x} & \vartheta \\ \mathbf{0}_{n_\theta,n_x} & \mathbf{0}_{n_\theta,n_\theta} \end{bmatrix}. \tag{5}$$

The term $\vartheta \in \mathbb{R}^{n_x \times n_\theta}$ serves as a mapping operator connecting relevant components to a nonlinear activation, defined as

$$\vartheta = \begin{bmatrix} 0 & \ldots & 0 \\ 1 & & 0 \\ \vdots & \ddots & \vdots \\ 0 & & 0 \\ 0 & \ldots & 1 \end{bmatrix}. \tag{6}$$

The discrete variant of **B** is expressed by

$$\mathbf{B}_\delta = \delta\mathbf{B}. \tag{7}$$

The connection matrix **C** remains identical for both continuous and discrete formulations, structured as

$$\mathbf{C} = \mathrm{diag}(\mathbf{\Pi}, \mathbf{0}_{n_\theta,n_\theta}). \tag{8}$$

The block $\mathbf{\Pi} \in \mathbb{R}^{n_x \times n_x}$ is actually an adjacency matrix (i.e., containing zeros and ones) defining the structure of local connections and consolidates incoming signals to compute mean states, subsequently passing them through an activation function. Its specific configuration is

$$\mathbf{\Pi} = \begin{bmatrix} 0 & 0 & \dots & 0 & 0 \\ c_{2,1} & 0 & & c_{2,n_x-1} & 0 \\ \vdots & & \ddots & & \vdots \\ 0 & 0 & & 0 & 0 \\ c_{n_x,1} & 0 & & c_{n_x,n_x-1} & 0 \end{bmatrix}. \quad (9)$$

For more detailed description of the neural mass model and its mathematical implementation, readers are encouraged to see [1].

### 2 Supplementary Tables

| Network | Brain structure | Network | Brain structure | Network | Brain structure |
|---|---|---|---|---|---|
| *DMN* | Parietal_Sup_L | *ECN* | Frontal_Sup_L | *SMN* | Precentral_L |
| | Parietal_Sup_R | | Frontal_Sup_R | | Precentral_R |
| | Parietal_Inf_L | | Frontal_Sup_Orb_L | | Frontal_Mid_L |
| | Parietal_Inf_R | | Frontal_Sup_Orb_R | | Frontal_Mid_R |
| | Temporal_Mid_L | | Frontal_Mid_L | | Supp_Motor_Area_L |
| | Temporal_Mid_R | | Frontal_Mid_R | | Supp_Motor_Area_R |
| | | | Frontal_Mid_Orb_L | | Postcentral_L |
| *DMNa* | Insula_L | | Frontal_Mid_Orb_R | | Postcentral_R |
| | Insula_R | | Frontal_Inf_Oper_L | | SupraMarginal_L |
| | Cingulum_Ant_L | | Frontal_Inf_Oper_R | | SupraMarginal_R |
| | Cingulum_Ant_R | | Frontal_Inf_Tri_L | | Paracentral_Lobule_L |
| | Temporal_Pole_Mid_L | | Frontal_Inf_Tri_R | | Paracentral_Lobule_R |
| | Temporal_Pole_Mid_R | | Frontal_Inf_Orb_L | | |
| | | | Frontal_Inf_Orb_R | *VN* | Calcarine_L |
| *DMNv* | Cingulum_Post_L | | Frontal_Sup_Medial_L | | Calcarine_R |
| | Cingulum_Post_R | | Frontal_Sup_Medial_R | | Cuneus_L |
| | Hippocampus_L | | Frontal_Med_Orb_L | | Cuneus_R |
| | Hippocampus_R | | Frontal_Med_Orb_R | | Lingual_L |
| | ParaHippocampal_L | | Rectus_L | | Lingual_R |
| | ParaHippocampal_R | | Rectus_R | | Occipital_Sup_L |
| | Fusiform_L | | Cingulum_Ant_L | | Occipital_Sup_R |
| | Fusiform_R | | Cingulum_Ant_R | | Occipital_Mid_L |
| | Angular_L | | Postcentral_L | | Occipital_Mid_R |
| | Angular_R | | Postcentral_R | | Occipital_Inf_L |
| | Precuneus_L | | Parietal_Sup_L | | Occipital_Inf_R |
| | Precuneus_R | | Parietal_Sup_R | | |
| | | | Parietal_Inf_L | | |
| *AN* | Insula_L | | Parietal_Inf_R | | |
| | Insula_R | | SupraMarginal_L | | |
| | Heschl_L | | SupraMarginal_R | | |
| | Heschl_R | | Angular_L | | |
| | Temporal_Sup_L | | Angular_R | | |
| | Temporal_Sup_R | | Precuneus_L | | |
| | | | Precuneus_R | | |

**Table S1** Brain structures in functional networks are shown in the format of Automated Anatomical Labeling atlas [2]. The specific brain structures are identified based on previous studies [3–5]. Seven functional networks: Default Mode Network (DMN), anterior Default Mode Network (DMNa), ventral Default Mode Network (DMNv), Auditory Network (AN), Executive Control Network (ECN), Sensorimotor Network (SMN), Visual Network (VN).

| Network | Brain structure | Network | Brain structure |
|---|---|---|---|
| *Prefrontal Network* | Frontal_Sup_L | *Posterior Parietal Network* | Postcentral_L |
| | Frontal_Sup_R | | Postcentral_R |
| | Frontal_Sup_Orb_L | | Parietal_Sup_L |
| | Frontal_Sup_Orb_R | | Parietal_Sup_R |
| | Frontal_Mid_L | | Parietal_Inf_L |
| | Frontal_Mid_R | | Parietal_Inf_R |
| | Frontal_Mid_Orb_L | | SupraMarginal_L |
| | Frontal_Mid_Orb_R | | SupraMarginal_R |
| | Frontal_Inf_Oper_L | | Angular_L |
| | Frontal_Inf_Oper_R | | Angular_R |
| | Frontal_Inf_Tri_L | | Precuneus_L |
| | Frontal_Inf_Tri_R | | Precuneus_R |
| | Frontal_Inf_Orb_L | | |
| | Frontal_Inf_Orb_R | | |
| | Frontal_Sup_Medial_L | | |
| | Frontal_Sup_Medial_R | | |
| | Frontal_Med_Orb_L | | |
| | Frontal_Med_Orb_R | | |
| | Rectus_L | | |
| | Rectus_R | | |

**Table S2** Brain structures in prefrontal network and posterior parietal network in the format of Automated Anatomical Labeling atlas. The specific brain structures are identified based on the previous study [6].

| | | Correlation Imaging | | | | Contrast Imaging | | | |
|---|---|---|---|---|---|---|---|---|---|
| Network | Brain structure | $\alpha_{pi}$ | $\alpha_{pe}$ | $\alpha_{ep}$ | $\mu$ | $\alpha_{pi}$ | $\alpha_{pe}$ | $\alpha_{ep}$ | $\mu$ |
| *DMN* | Parietal_Sup_L | | | 3.9389 | 3.7324 | | | -8.4166 | |
| | Parietal_Sup_R | | | 3.4843 | | | | | |
| | Parietal_Inf_L | | | | | | | | |
| | Parietal_Inf_R | | | 3.4857 | | | | | |
| | Temporal_Mid_L | | | | | | | | |
| | Temporal_Mid_R | | | | | | | | -3.4651 |
| *DMNa* | Insula_L | | | | | | | | |
| | Insula_R | | | | | | | | -3.3862 |
| | Cingulum_Ant_L | | | | | | -4.1818 | | |
| | Cingulum_Ant_R | | | | | | | | |
| | Temporal_Pole_Mid_L | 3.7224 | 3.7168 | | | -4.0599 | | | |
| | Temporal_Pole_Mid_R | | | | | | -4.8823 | | |
| *DMNv* | Cingulum_Post_L | | 4.0126 | | | | -4.1181 | | |
| | Cingulum_Post_R | | | 3.6354 | | | | | |
| | Hippocampus_L | | | | | | | | |
| | Hippocampus_R | | | | | | | | |
| | ParaHippocampal_L | | | | | | | | -3.4271 |
| | ParaHippocampal_R | | | | | | | | -4.2258 |
| | Fusiform_L | | | | | | | | |
| | Fusiform_R | | | | | | -4.6501 | | |
| | Angular_L | | | 3.7478 | 3.5735 | | | -9.3609 | |
| | Angular_R | | | | | | | | |
| | Precuneus_L | | | 3.2482 | | | -3.904 | | |
| | Precuneus_R | | 3.4036 | 3.5353 | | | | | |
| *SMN* | Precentral_L | | | | | | | | |
| | Precentral_R | | | | | | | | |
| | Frontal_Mid_L | | | | | | | | |
| | Frontal_Mid_R | | | | | | | | |
| | Supp_Motor_Area_L | | | | | | | | |
| | Supp_Motor_Area_R | | | | | | | | |
| | Postcentral_L | | | | | | | | |
| | Postcentral_R | | | | | | | | |
| | SupraMarginal_L | | | | | | | | |
| | SupraMarginal_R | | | | 3.6124 | | | | |
| | Paracentral_Lobule_L | | | | | | | | |
| | Paracentral_Lobule_R | | | 3.2410 | | | | | |
| *VN* | Calcarine_L | | | 3.2163 | | | | -7.0919 | |
| | Calcarine_R | | 3.4504 | 3.8012 | 3.5762 | | | -7.3398 | |
| | Cuneus_L | | 3.5177 | 3.4961 | 3.3911 | | | -9.2590 | -3.7809 |
| | Cuneus_R | | | 3.5109 | | | | | |
| | Lingual_L | | | 3.9112 | 4.1726 | | | -7.1325 | -3.8016 |
| | Lingual_R | | 3.8917 | | | | | | -3.4081 |
| | Occipital_Sup_L | | 3.8459 | | | | | -7.0552 | |
| | Occipital_Sup_R | | | 3.7282 | | | | | |
| | Occipital_Mid_L | | | 4.0056 | | | | | |
| | Occipital_Mid_R | | | | | | | -9.0837 | |
| | Occipital_Inf_L | | | 3.5362 | | | | -7.8248 | |
| | Occipital_Inf_R | | | 3.3933 | | | | -8.7592 | |
| *AN* | Insula_L | | | | | | | | |
| | Insula_R | | | | | | | | -3.3862 |
| | Heschl_L | | | | | | | | -3.4770 |
| | Heschl_R | | | | | | | | |
| | Temporal_Sup_L | | | | | | | | |
| | Temporal_Sup_R | | | | | | | | |
| *ECN* | Frontal_Sup_L | | | | | | | | -3.4626 |
| | Frontal_Sup_R | | | | | | | | -3.9891 |
| | Frontal_Sup_Orb_L | | | | | -4.7673 | | | |
| | Frontal_Sup_Orb_R | | | | | | | | |
| | Frontal_Mid_L | | | | | | | | |
| | Frontal_Mid_R | | | | | | | | |
| | Frontal_Mid_Orb_L | 4.2717 | | | | -4.7284 | | | |
| | Frontal_Mid_Orb_R | | | | | | | | |
| | Frontal_Inf_Oper_L | | | | | | | | -4.3361 |
| | Frontal_Inf_Oper_R | | | | | | | | |
| | Frontal_Inf_Tri_L | | | | | | | | |
| | Frontal_Inf_Tri_R | | | | | | | | |
| | Frontal_Inf_Orb_L | | | | | -3.7773 | | | |
| | Frontal_Inf_Orb_R | | | | | | | | |
| | Frontal_Sup_Medial_L | | | | | | | | |
| | Frontal_Sup_Medial_R | | | | | | | | -3.5087 |
| | Frontal_Med_Orb_L | | 3.4132 | | | -4.1911 | | | |
| | Frontal_Med_Orb_R | | 3.4348 | | | | | | |
| | Rectus_L | | | | | | -5.6424 | | |
| | Rectus_R | | | | | | | | |
| | Cingulum_Ant_L | | | | | | -4.1818 | | |
| | Cingulum_Ant_R | | | | | | | | |
| | Postcentral_L | | | | | | | | |
| | Postcentral_R | | | | | | | | |
| | Parietal_Sup_L | | | 3.9389 | 3.7324 | | | -8.4166 | |
| | Parietal_Sup_R | | | 3.4843 | | | | | |
| | Parietal_Inf_L | | | | | | | | |
| | Parietal_Inf_R | | | 3.4857 | | | | | |
| | SupraMarginal_L | | | | | | | | |
| | SupraMarginal_R | | | | 3.6124 | | | | |
| | Angular_L | | | 3.7478 | 3.5735 | | | -9.3609 | |
| | Angular_R | | | | | | | | |
| | Precuneus_L | | | 3.2482 | | | -3.9040 | | |
| | Precuneus_R | | 3.4036 | 3.5353 | | | | | |

**Table S3** Functional networks, and the corresponding brain structures and group-level t-statistics for regional neurophysiological variables in correlation imaging and contrast imaging. Multiple comparisons permutation tests were used (see "Methods") [7].

### 3 Supplementary Figures

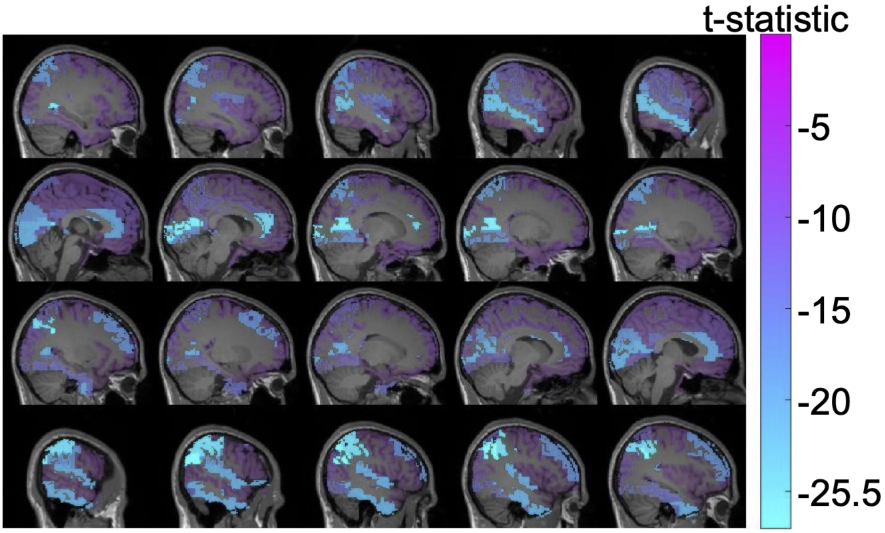


**Fig. S1** The cortical regions where the membrane potential of the pyramidal population exhibited significant differences before and after Xenon equilibrated are presented. Significant regions were identified by t-statistics derived from the multiple comparisons permutation test at the significance level $\alpha = 0.05$.

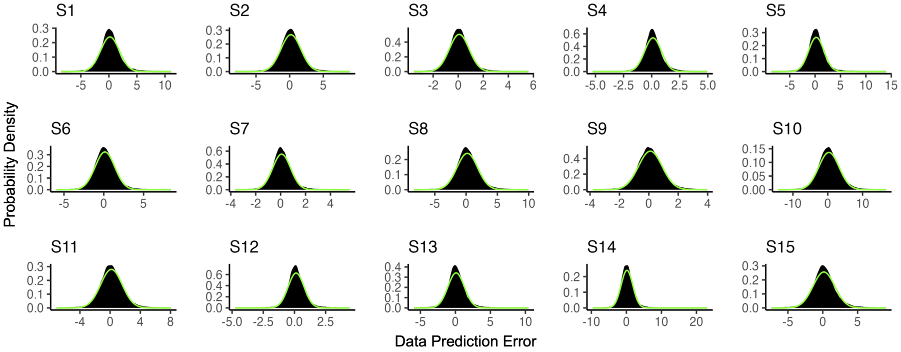


**Fig. S2** Neural mass models were forward simulated using parameter estimates. The data prediction error is the difference between the source-level MEG time series and the forward simulation. The data prediction errors for all 15 subjects are depicted as probability density functions (black histograms), with normal probability density functions having equivalent mean and standard deviation (green curves).
